# Sequential Measurements of Catalytic Activities of Multi-Drug-Resistance Transporters and Cytochrome P450 Enzymes by Cytometry of Reaction Rate Constant

**DOI:** 10.1101/2020.10.15.340976

**Authors:** Vasilij Koshkin, Mariana Bleker de Oliveira, Sven Kochmann, Chun Peng, Sergey N. Krylov

## Abstract

Cytometry of reaction rate constant (CRRC) is an accurate and robust approach to characterize cell-population heterogeneity using rate constants of cellular processes for which kinetic mechanisms are known. We work on a CRRC-based method to develop predictors of tumor chemoresistance driven by two processes: drug extrusion by multi-drug-resistance (MDR) transporters and drug inactivation by cytochrome-P450 enzymes (CYP). Each of the two possess is studied with its specific substrate and the process activity is characterized by a corresponding unimolecular rate constant. Due to the incompatibility of MDR and CYP assays, MDR and CYP activities may be difficult to measure simultaneously suggesting that they may need to be measured sequentially. The sequential measurements may also impose a problem: the results of the second assay may be affected by artifacts exerted by the first assay. The goal of this work was to understand whether the cells have a memory of the first assay that significantly affects the results of the second assay. To achieve this goal, we compared CRRC results for two orders of sequential measurements: the MDR→CYP order in which MDR activity is measured before CYP activity and the CYP→MDR order in which CYP activity is measured before MDR activity. It was found that the results of the CYP assay were similar in both orders; on the contrary, the results of the MDR assay were significantly different. Our findings suggest that MDR and CYP activity can be studied sequentially provided that MDR activity is measured first and CYP activity second.

## Introduction

Cancer resistance to primary chemotherapy is mainly caused by a small population of tumor cells with increased drug resistance. Among major mechanisms of cellular resistance to drugs are accelerated drug extrusion from cells by multi-drug-resistance (MDR) transporters and accelerate drug inactivation by intracellular enzymes from the cytochrome-P450 (CYP) family.^1^ Such drug-resistant cells have a higher probability of surviving primary chemotherapy and giving rise to a drug-resistant tumor. Therefore, the relative size of the drug-resistant cell population is viewed as a marker of tumor chemoresistance.^2^ Finding a chemoresistance predictor based on the size of the drug-resistant cell population requires accurate determination of this size *via* measuring MDR and CYP activities at the single-cell level, *i.e.*, by cytometry using fluorescent (for MDR) and fluorogenic (for CYP) substrates.^3^ Flow cytometry is not accurate and not robust for such measurements.^4^ Cytometry of reaction rate constant (CRRC), which is based on time-lapse fluorescence microscopy, is, in contrast, accurate and robust. Being such, CRRC is a highly-promising platform for development of chemoresistance predictors.^5^ CRRC monitors kinetics of cellular processes in individual cells and uses a reaction rate constant of the process (with known kinetic mechanism) to characterize the heterogeneity of the cell population. CRRC was comprehensively evaluated in measurements of MDR activity during the past decade.^5,6^ In contrast, using CRRC for studies of CYP activity is still in its infancy.

MDR and CYP activities can technically be measured simultaneously (in two different optical channels), but there are biochemical interferences that can affect the accuracy of simultaneous measurements and which force us to consider sequential measurements. However, even in sequential measurements, the cellular changes, which the substrate and inhibitors of the first process introduce, may affect the results of measurements of the second process. In essence, the sequential analysis is only feasible if the sequence order of the measurements does not influence the rate constants. In this work, we assessed if a method based on CRRC for the sequential investigation of MDR and CYP activities was robust towards the change of the sequence order. Accordingly, we studied two sequence orders: the MDR→CYP order in which MDR activity is measured before CYP activity and the CYP→MDR order in which measurements of MDR activity follow that of CYP activity. The results were presented as kinetic histograms “number of cells vs rate constant” and scatter plots “MDR rate constant (*k*_MDR_) vs CYP rate constant (*k*_CYP_)”. We found that the CYP activity was similar for the MDR→CYP and CYP→MDR orders, while the MDR activity was significantly higher for the MDR→CYP order. Our results strongly suggest that the sequential analysis of the two activities by CRRC is feasible if MDR activity is measured first followed by assessing the CYP activity.

## Materials and methods

### Reagents

All reagents were obtained from Sigma-Aldrich (Oakville, Ontario, Canada), Fluka AG (Buchs, Switzerland), and BDH Chemicals Ltd. (Poole, England).

### Instrumentation

All measurements were performed with a Leica DMi8 fluorescence microscope.

### Cell culture

A2780 ovarian cancer cells were grown in Eagle’s Minimum Essential Medium (EMEM) and supplemented with 100 IU/mL penicillin, 100 μg/mL streptomycin, and 10% fetal bovine serum in a humidified atmosphere of 5% CO2 at 37 °C. The moderate content of CYP in these cells^7^ was increased by addition of phenobarbital (0.2 mM) for 24 h.^8^

### Kinetic MDR assay

MDR efflux was monitored by the cell extrusion of fluorescein that resulted in a fluorescence decrease inside the cells. The detailed imaging procedure is described elsewhere.^5^ Briefly, the cell plate was filled with Hank’s Balanced Salt Solution (HBSS) medium. Fluorescein (substrate, 1.5 µM) and glibenclamide (inhibitor, 10 μM) were added to the cells for 30 min to load fluorescein into the cells. The loading was stopped by removing the extracellular substrate by carefully replacing the cell support medium with a fresh Krebs-Ringer-Bicarbonate (KRB) buffer supplemented with glibenclamide (10 μM) but without fluorescein. This replacement initiated passive substrate leakage through the membrane that was monitored for 10–15 min. Then, the MDR-mediated fluorescein efflux was initiated by replacing the inhibitor-containing cell support medium with pure HBSS, *i.e.* without any substrate or inhibitor. The decreasing fluorescence intensity caused by the MDR transport was monitored for 60–90 min (efflux was visually completed by the end of measurements). All cells within the field of view were imaged with 3 min intervals.

### Kinetic CYP assay

CYP activity was monitored by the o-dealkylation of pentoxyresorufin (fluorogenic substrate) to resorufin (fluorescent product) over time that resulted in a fluorescence increase inside the cells. The detailed imaging procedure is described elsewhere.^9^ Briefly, the cell plate was filled with the HBSS medium. The medium was supplemented with probenecid (1 mM) and dicoumarol (25 µM), to inhibit the efflux of resorufin and the transformation of resorufin to dihydroresorufin, respectively. Next, pentoxyresorufin (5 µM) was added to the cells. The increasing fluorescence was monitored for 15–20 min; all cells in the field of view were imaged with 1 min intervals. Finally, identification of individual cells was performed with propidium-iodide and saponin staining.

### Sequential assays

Kinetic MDR and CYP assays were performed in two sequence orders: MDR→CYP and CYP→MDR. The cell plate was washed and filled with the respective assay medium for the subsequent measurement after the previous assay.

### Extraction and analysis of kinetic traces

Regions of interest (ROIs) representing individual cells were analysed through stacks of the time-lapse images acquired during MDR and CYP measurements using *ImageJ software*.^10^ Kinetic traces of each individual cell were extracted by using the mean fluorescence signal of the respective cell over time. Values of *k*_MDR_ were determined by using kinetic traces of fluorescein efflux in each individual cell as described previously.^5^ Values of *k*_CYP_ were determined similarly by fitting kinetic traces of resorufin formation in each individual cell using first-order kinetics.^11^ All fitting procedures were performed using *OriginPro software*.

### Cell population analysis

MDR and CYP activities withing cell populations were characterized by histograms representing distributions of *k*_MDR_ and *k*_CYP_ in single cells across cell populations. Histograms were plotted in *OriginPro software* using the *Automatic Binning mode* and characterized by the median and interquartile range (middle 50%) as described previously.^5^ Statistical parameters were determined using *OriginPro’s Descriptive Statistics* tool. Clusters were analysed by a self-developed Python program and the *k*-means algorithm (with *k* = 2 clusters).

### Availability of data

Raw data, evaluation files, and the Python program for cluster analysis can be found in the supplementary information.

## Results and Discussion

### Incompatibility of MDR and CYP assays

Simultaneous measurement of MDR and CYP activities may not be possible due to biochemical incompatibility: resorufin, which is produced and used as the fluorescent analyte during the CYP assay, will be removed from the cells by MDR transport.^12^ If fast relative to resorufin production, this removal will decrease the measured fluorescence signal *inside* the cell significantly leading to apparently lower CYP activity. Thus, the CYP assay requires the addition of an MDR inhibitor (here: probenecid) to suppress MDR activity and prevent the underestimation of the CYP activity. At the same time, the suppressed MDR activity makes it impossible to assess the native MDR activity. Conclusively, MDR and CYP assays are biochemically incompatible in general and have to be performed sequentially.

### Requirements of an optimal sequential assay

Any assay in the sequence should not change the physiological state of the analysed cells significantly and permanently not to affect the next assay. The effects of a preceding assay (*e.g.* inhibition) on the results of the subsequent assays should be negligible. Otherwise, the subsequently determined rate constants become dependent on the previous assays. Consequently, the order of assays in the sequential measurement must be chosen so that the measurements do not interfere, and the results of the subsequent assays do not depend on the presence of the previous assays.

### Sequential assays involving MDR and CYP

Here, we assess if the sequential measurement of MDR and CYP activities is robust towards the change of sequence order. Accordingly, we studied two sequence orders: the MDR→CYP order, in which MDR activity is measured before CYP activity, and the CYP→MDR order, in which measurements of MDR activity follow that of CYP activity. We present the CRRC results as kinetic histograms “number of cells vs rate constant” (Figure 1) and scatter plots “MDR rate constant (*k*_MDR_) vs CYP rate constant (*k*_CYP_)” (Figure 2).

**Figure 1.**
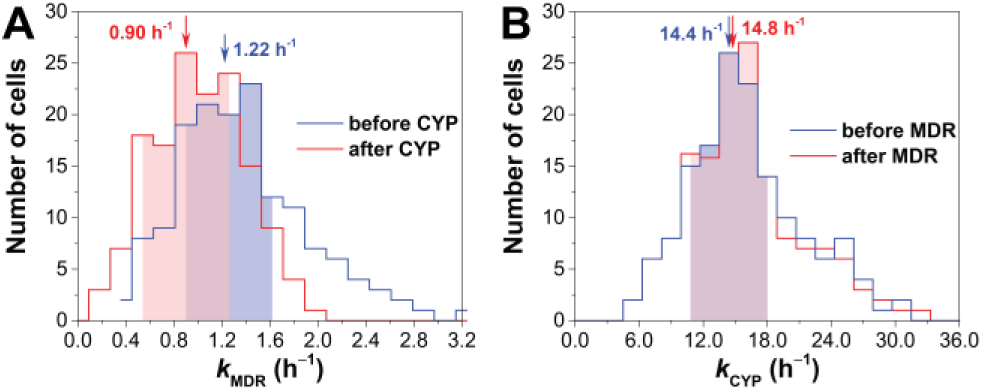
Histograms of MDR (A) and CYP (B) activities in sequential MDR and CYP assays. While the CYP activity is not influenced significantly by changing the sequence order, the MDR activity is shifted to lower values when switching from the MDR→CYP order to the CYP→MDR order.

**Figure 2.**
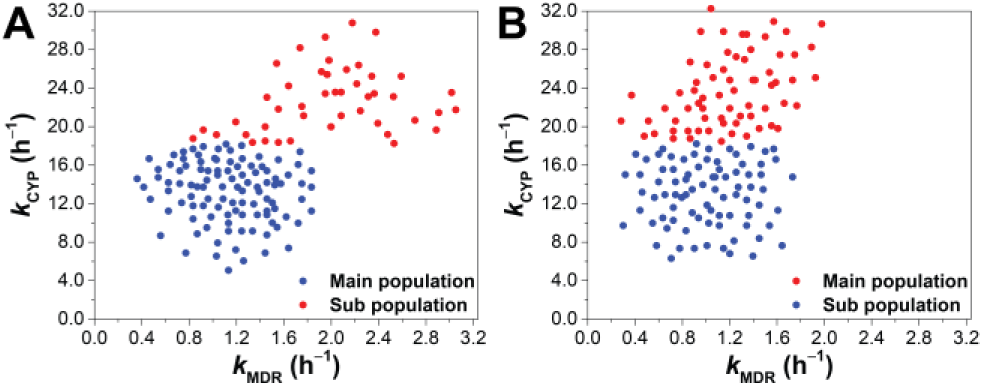
Bivariate distributions of MDR and CYP activities for the MDR→CYP (A) and CYP→MDR (B) assay sequence orders. The clusters were analysed by *k*-means.

We found that the MDR activity was significantly higher for the MDR→CYP order than for the CYP→MDR order (Figure 1A). Performing the CYP assay first affected the MDR assay result by shifting both the median of the distribution (from 1.22 h^−1^ to 0.90 h^−1^) and the interquartile range (from [0.90‒1.62 h^−1^] to [0.61‒1.19 h^−1^]). In contrast, the CYP activity was similar for the MDR→CYP and CYP→MDR orders (Figure 1B); neither the median of the distribution (14.8 h^−1^ vs h^−1^) nor the interquartile range ([10.8‒18.0 h^−1^] vs. [11.2‒17.6 h^−1^]) changed significantly (changes < 5%). These results strongly suggest that the sequential analysis of the two activities by CRRC is feasible for the MDR→CYP sequence order only.

### Cluster analysis

Bivariate plots of *k*_CYP_ vs *k*_MDR_ can provide additional information on the relationship between MDR and CYP activities (Figure 2). The *k*_CYP_ vs *k*_MDR_ plot of the data obtained for the MDR→CYP order (Figure 2A) clearly shows two separate clusters; this suggests the existence of a separate cell subpopulation with elevated activities of both MDR and CYP. Such cell subpopulations of cancer cells (here: A2780 ovarian cancer cells) may represent cancer stem (tumor-initiating) cells. This distinct subpopulation in Figure 2A is not obvious in both respective histograms of MDR and CYP activities (Figure 1). However, this subpopulation is less distinctively visible in data obtained for the CYP→MDR order (Figure 2B). The lack of a distinct separator between the two clusters is due to the reduced and underestimated activity in the MDR assay performed after the CYP assay. This cluster analysis confirms that the sequential measurement of MDR and CYP activities is only possible for the MDR→CYP sequence order, and that assessing of multiple drug removal pathways leads to better assessment of the heterogeneity of cell populations.

## Conclusion

In this work, we assessed if a method based on CRRC for the sequential investigation of MDR and CYP activities was robust towards the change in assay sequence order. Accordingly, we studied two sequence orders: MDR→CYP and CYP→MDR. We found that the CYP activity was similar for the MDR→CYP and CYP→MDR orders, while the MDR activity was significantly higher for the MDR→CYP case. The results show that the accurate assessment of the cellular drug extrusion by MDR transporters and drug inactivation by CYP enzymes requires that the CRRC extrusion assay be performed first. Otherwise, drug extrusion (MDR) activity is underestimated due to the residual toxic effects of metabolic inhibitors present in the drug inactivation (CYP) assay. Conclusively, the sequential analysis of both MDR and CYP activity by CRRC is feasible in for the MDR→CYP assay sequence order.

## Supporting information

Raw data, evaluation data, and cluster analysis program (Python)

