## Supplementary material for "Sequential Measurements of Catalytic Activities of Multi-Drug-Resistance Transporters and Cytochrome P450 Enzymes by Cytometry of Reaction Rate Constant": Raw data, evaluation data, and cluster analysis program (Python): clusteranalysis.py

```
#!/usr/bin/env python3
# -*- coding: utf-8 -*-

# Copyright (C) 2020 by Sven Kochmann

# This program is free software: you can redistribute it and/or modify
# it under the terms of the GNU General Public License as published by
# the  Free Software Foundation,  either version 3  of the License, or
# (at your option) any later version.

# This program  is distributed  in the hope  that it will  be  useful,
# but  WITHOUT  ANY  WARRANTY;  without even  the implied warranty  of
# MERCHANTABILITY  or  FITNESS  FOR  A  PARTICULAR  PURPOSE.  See  the
# GNU General Public License for more details.

# You  should  have  received a copy of the GNU General Public License
# along with this program. If not, see <http://www.gnu.org/licenses/>.

# This script takes an X-Y data file and tries to sort all data points
# into two clusters by k-means (see scipy documentation).

import csv
import sys
from matplotlib import pyplot as plt
import scipy.cluster.vq

# Data file to operate on
if len(sys.argv) != 2:
    print("Syntax: clusteranalysis.py [datafile]")
    sys.exit(0)

# Reading the given data file
data = []
with open(sys.argv[1], newline='') as csvfile:
    reader = csv.reader(csvfile, delimiter=';')

    for row in reader:
        data.append([float(row[0]), float(row[1])])

# Computate the k-Means with k = 2 (assuming there are two clusters in the data)
centroids,_ = scipy.cluster.vq.kmeans(data,2)
idx,_ = scipy.cluster.vq.vq(data,centroids)

print("Centeroids:")
print(" Cluster 1:", centroids[:,0][0],centroids[:,1][0])
print(" Cluster 2:", centroids[:,0][1],centroids[:,1][1])
print("")

print("Data:")
for i, item in enumerate(data):
    if idx[i] == 0:
        plt.plot(item[0], item[1], 'ob')
        print(item[0], item[1], '', sep=';')
    else:
        plt.plot(item[0], item[1], 'or')
        print(item[0], '', item[1], sep=';')

# Plot the whole clusters as a check
plt.plot(centroids[:,0],centroids[:,1],'sg',markersize=8)
plt.show()
```
